# Establishing thresholds for cytokine storm and defining their relationship to disease severity in respiratory viral infections

**DOI:** 10.1101/2023.07.06.548022

**Authors:** Aisha Souquette, Jeremy Chase Crawford, Joshua Wolf, Ashley Blair, Mona Agrawal, Laura Boywid, Kristin McNair, Nicholas D. Hysmith, Chloe Hundman, Azadeh Bahadoran, Sandra R. Arnold, SJTRC Study Team, Heather S. Smallwood, Amanda Green, Paul G. Thomas

## Abstract

Previous studies have identified cytokines associated with respiratory virus infection illness outcome. However, few studies have included comprehensive cytokine panels, longitudinal analyses, and/or simultaneous assessment across the severity spectrum. This, coupled with subjective definitions of cytokine storm syndrome (CSS), have contributed to inconsistent findings of cytokine signatures, particularly with COVID severity. Here, we measured 38 plasma cytokines and compared profiles in healthy, SARS-CoV-2 infected, and multisystem inflammatory syndrome in children (MIS-C) patients (n = 169). Infected patients spanned the severity spectrum and were classified as Asymptomatic, Mild, Moderate or Severe. Our results showed acute cytokine profiles and longitudinal dynamics of IL1Ra, IL10, MIP1b, and IP10 can differentiate COVID severity groups. Only 4% of acutely infected patients exhibited hypercytokinemia. Of these subjects, 3 were Mild, 3 Moderate, and 1 Severe, highlighting the lack of association between CSS and COVID severity. Additionally, we identified IL1Ra and TNFa as potential biomarkers for patients at high risk for long COVID. Lastly, we compare hypercytokinemia profiles across COVID and influenza patients and show distinct elevated cytokine signatures, wherein influenza induces the most elevated cytokine profile. Together, these results identify key analytes that, if obtained at early time points, can predict COVID illness outcome and/or risk of complications, and provide novel insight for improving the conceptual framework of hypercytokinemia, wherein CSS is a subgroup that requires concomitant severe clinical manifestations, and including a list of cytokines that can distinguish between subtypes of hypercytokinemia.

## Introduction

Cytokines are an ideal biomarker, which can be quickly measured with a small volume of easily obtained samples, and are easy therapeutic targets, with treatments already available. Although numerous cytokine correlates of COVID severity have been identified (Leisman et al. 2020), there have been few longitudinal studies with simultaneous assessment across the spectrum of disease severity. This makes it difficult to determine how many severity profiles exist and the number of therapeutic approaches required to optimally treat patients. Moreover, the role of cytokine storm syndrome (CSS) as a driver of severe disease remains poorly understood, in part due to subjective definitions of CSS and differences in comparator groups across studies (Mudd et al. 2020; Fajgenbaum and June 2020). Additionally, few studies assess more than 6 analytes, and fewer still control for type I error in their statistical analyses of multiple analytes.

Here, we measure the levels of 38 plasma cytokines from 169 COVID patients who exhibited a broad spectrum of disease severity, characterize analytes uniquely associated with illness outcomes, and outline a strict statistical definition for COVID associated hypercytokinemia (CAH), a proposed subset of hypercytokinemia that occurs during acute infection and is distinct from CSS. Our results demonstrate that cytokine profiles and dynamics vary as a function of disease severity. CAH, MIS-C, and non-hypercytokinemia profiles can be distinguished by IL1b, GCSF, and IFNg. CAH occurred in only 4% of patients from across the illness-severity spectrum, making MIS-C the only representation of true CSS associated with severe disease. Additionally, we identify IL1Ra and TNFa as independent, putative biomarkers for individuals at high risk for long COVID (prolonged, post-acute COVID symptoms). Lastly, we show influenza virus infection induces the most elevated cytokine profile with hypercytokinemia signatures that are distinct from those during CAH or MIS-C. These results provide valuable insight into the immunological signatures associated with the severity of respiratory virus infection, which can be further utilized in clinical screening, and suggest multiple therapeutic options may be required to optimally treat patients.

## Results

### Clinical and demographic risk factors of severe COVID

The cohort was composed of two prospective, longitudinal study populations from Memphis, Tennessee: the SJTRC and the CIVIC-19 studies (Minervina et al. 2022). A total of 169 subjects were included who either tested positive for SARS-CoV-2 by RT-PCR (n = 164) or were diagnosed with MIS-C (n = 5). Reference time points include day(s) since PCR positive test, day(s) since symptom onset, and study day (time since first sample collection, median 7 and range -3 to 29 days post symptom onset for the most acute time point).

Demographic and clinical information was collected from electronic medical records, as available, and patient questionnaires (**Supp Table 1**). Participants were 7-89 years old (median age = 40), 69% female, 18.9% Black/African American, 59.8% White/Caucasian, and 2.4% Hispanic. Prevalent comorbidities included obesity (n = 56, 33.1%), heart disease (n = 30, 17.8%), chronic respiratory disease (n = 26, 15.4%), diabetes (n = 21, 12.4%), and current or past smoker (n = 25, 14.8%). Subjects were classified as Asymptomatic (n = 8, 4.7%), Mild (n = 117, 69.2%), Moderate (n = 22, 13.0%), or Severe (n = 17, 10.1%), as described in Methods. Consistent with prior work, analysis of demographic and comorbidity prevalence among severity groups showed diabetes (p = 0.0005) and hypertension (p = 0.003) are risk factors for severe COVID (**Fig 1**) (Gallo Marin et al. 2021).Therapeutic agents were given at the discretion of the participants’ treating clinicians. Hospitalized participants received a variety of treatments, most commonly high dose steroids, remdesivir, and/or antibiotics, including half of Moderates, all Severe, and all MIS-C. Aside from cytokine dynamics, all analyses were restricted to the most acute sample available. Of note, observed cytokine elevations occurred even after administration of anti-inflammatory agents.

**Figure 1.**
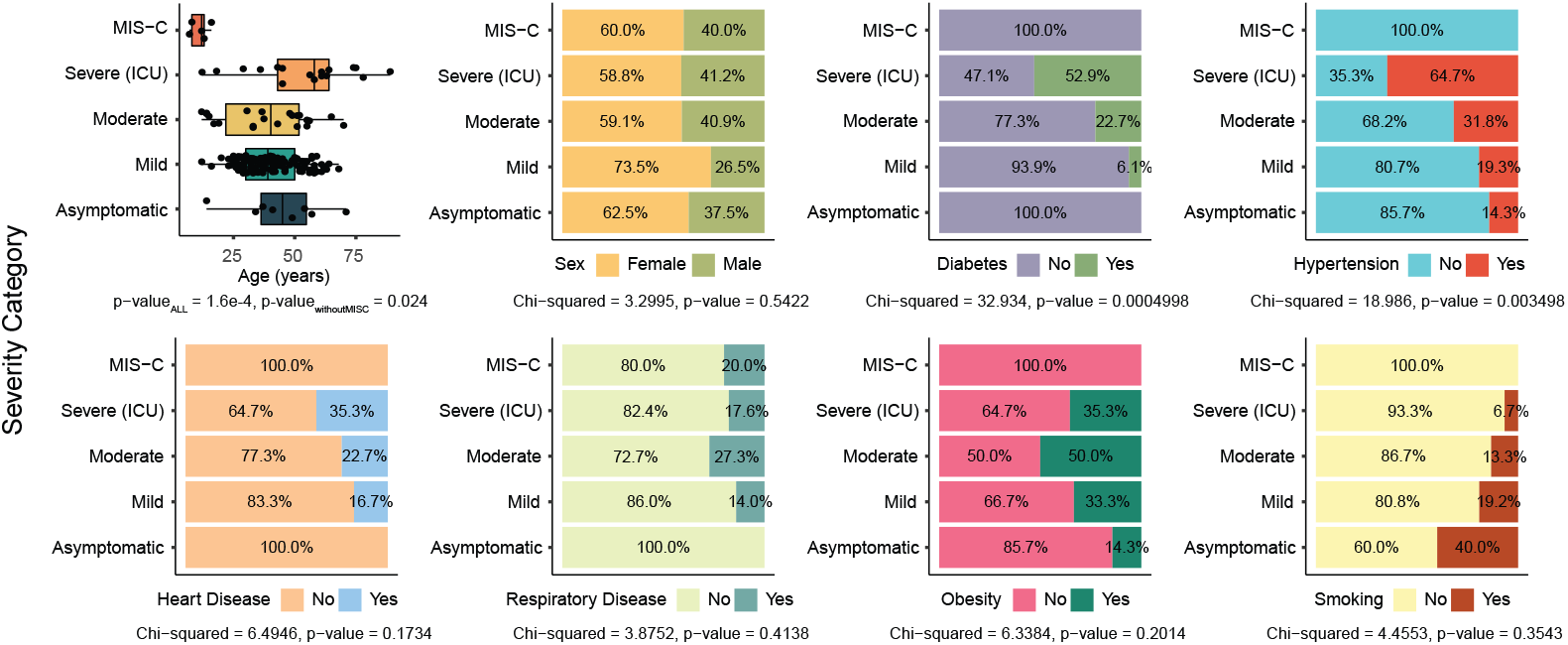
Clinical and demographic associations with COVID severity. Box and bar plots show age, sex, and prevalence of comorbidities across COVID severity groups. Boxplot elements are as follows: center line, median; box limits, lower and upper quartiles; whiskers, 1.5x interquartile range; dots, individual data points. Presented p-values are from Kruskal-Wallis or chi-squared tests.

### Cytokine profiles across the COVID severity spectrum

We measured the levels of 38 plasma cytokines and, based on our prior work, classified a subject as having “COVID associated hypercytokinemia” (CAH) if 30% or more of their cytokines were >2 standard deviations (SDs) from the acute infected mean cytokine level (Mudd et al. 2020). To prevent skewing, this calculation was determined using all patients except MIS-C due to its defining hyperinflammatory pathophysiology. Utilizing this metric, 7 of 164 subjects (4.27%) exhibited CAH, with 31.6-60.5% of their cytokines exceeding the acute infected mean by 2-6.8 SDs.

Principal component analysis (PCA) showed generally distinct cytokine profiles associated with COVID severity. The CAH cluster was closest to and partially overlapped with the MIS-C cluster, suggesting our quantitative approach appropriately separated a hypercytokinemia profile (**Fig 2A**). The groups furthest from each other, representing the most disparate profiles, were Asymptomatic and MIS-C. Moderate and Severe clusters partially overlapped and were positioned 3 times closer to Asymptomatic-Mild clusters than CAH-MIS-C clusters, highlighting the relative rarity of extreme hypercytokinemia in severe COVID. Differences in cytokine abundance across COVID severity groups were also observed among individual cytokines, with two prominent patterns (**Fig 2B** and **Supp Fig 1**): 1) A progressive increase in cytokine levels, as severity increased from Asymptomatic to Severe. This pattern included IL1Ra, IL8, IL10, and IL15, among others. 2) A progressive increase in cytokine levels from Asymptomatic to Moderate, but a decrease for Severe. Examples of this pattern include IL12p40, IFNg, IL1b, and TNFb, among others. Merging CAH subjects into their original severity group categorization did not change cytokine patterns, but generally increased variance (**Supp Fig 2-3**).

**Figure 2.**
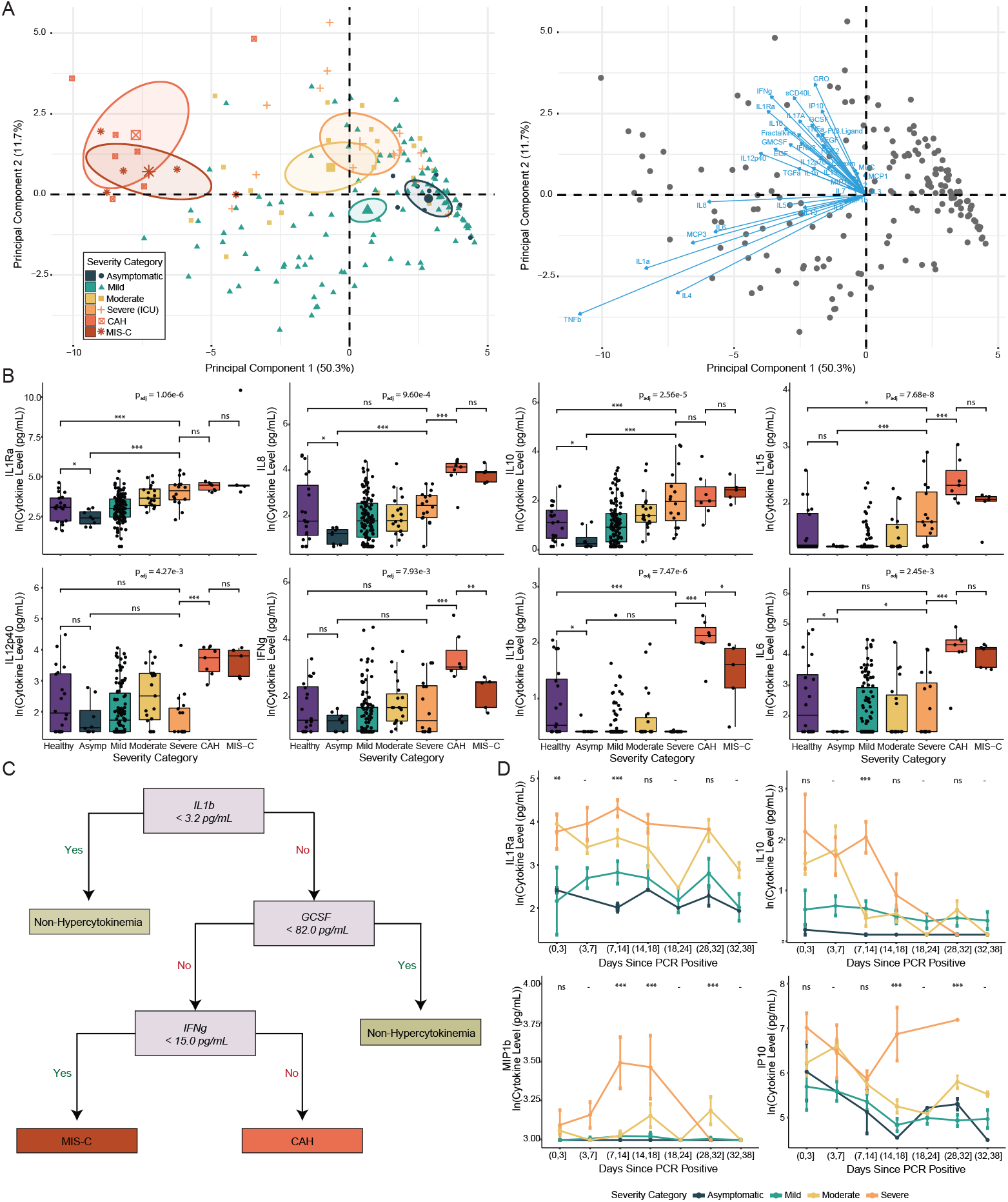
Unique cytokine profiles and dynamics across the COVID severity spectrum. (A) Principal component analysis of 38 cytokines. (B) Box plots of acute cytokine levels by COVID severity group. Boxplot elements are as follows: center line, median; box limits, lower and upper quartiles; whiskers, 1.5x interquartile range; dots, individual data points.Written p-value is from a Kruskal-Wallis test for all groups, except CAH. (C) Classification tree analysis to distinguish non-hypercytokinemia, CAH, and MIS-C immune profiles. (D) Cytokine dynamics by COVID severity group (n_Total_ = 151, n_Unique_ = 60). Points represent the mean and lines represent standard error. Stars or “ns” represent Mann-Whitney U (B) or Kruskal-Wallis (D) tests as follows: raw p-value < 0.05 and * p_adj_ < 0.2, ** p_adj_ < 0.1, *** p_adj_ < 0.05, “ns” = not significant.

Patients with severe disease significantly differed from healthy controls by 9 analytes, 2 decreased (IL13, IL1b) and 7 increased (GCSF, GRO, IL10, IL15, sCD40L, IL1Ra, and IP10) (**Fig 2B, Supp Fig 1**). Interestingly, Asymptomatic patients exhibited lower TGFa, MIP1b, IL6, IL8, IL10, and IL1Ra compared to both Healthy and Severe groups (**Fig 2B, Supp Fig 1**). Although CAH and MIS-C were both hypercytokinemia profiles (**Fig 2A-B, Supp Fig 4**), there were select differences between them, such as IFNg and IL1b (**Fig 2B**), suggesting there may be unique pathophysiologies that warrant further investigation. These results coincided with a classification tree analysis, which showed that IL1b, GCSF, and IFNg were sufficient to distinguish between CAH, MIS-C, and non-hypercytokinemia subjects with an overall misclassification rate of 1.6% (**Fig 2C)**. Data from our prior independent study of cytokine profiles in hospitalized adult COVID patients validated these results, IL1b and GCSF accurately classified CAH and non-hypercytokinemia subjects with a misclassification rate of 2.53% (2 out of 79) (Mudd et al. 2020). Of note, these data were collected using a different assay and in a different lab, highlighting the robustness of these classifiers.

Further delineation between Asymptomatic and Severe COVID is evident in the dynamics of IL1Ra, IL10, MIP1b, and IP10 (**Fig 2D** and **Supp Fig 5**). At early time points, high levels of IL1Ra clearly distinguished Moderate-Severe from Asymptomatic-Mild cases. At day 7 to 14, an early peak of IL1Ra distinguished Mild from Asymptomatic cases, and Severe cases exhibited uniquely high levels of IL10 and MIP1b. While all severity groups had similar IP10 levels at early time points, Severe patients diverged after day 7-14, such that IP10 levels progressively increased over time. For all other severity groups, levels of IP10 decreased with time, leveling off around day 14-18.

Long COVID is a condition in which patients experience continuous or relapsing, prolonged symptom duration for several weeks to months (Michelen et al. 2021). Here, “long COVID” was defined as any COVID symptoms persisting for greater than 4 weeks. Of the 145 patients with available records, 37 (26%) experienced long COVID. Of those, 23 were Mild, 6 Moderate, and 8 Severe (20%, 40%, and 80% of their severity categories, respectively) in the first 28 days of illness (detailed in Methods) (**Fig 3A**).

**Figure 3.**
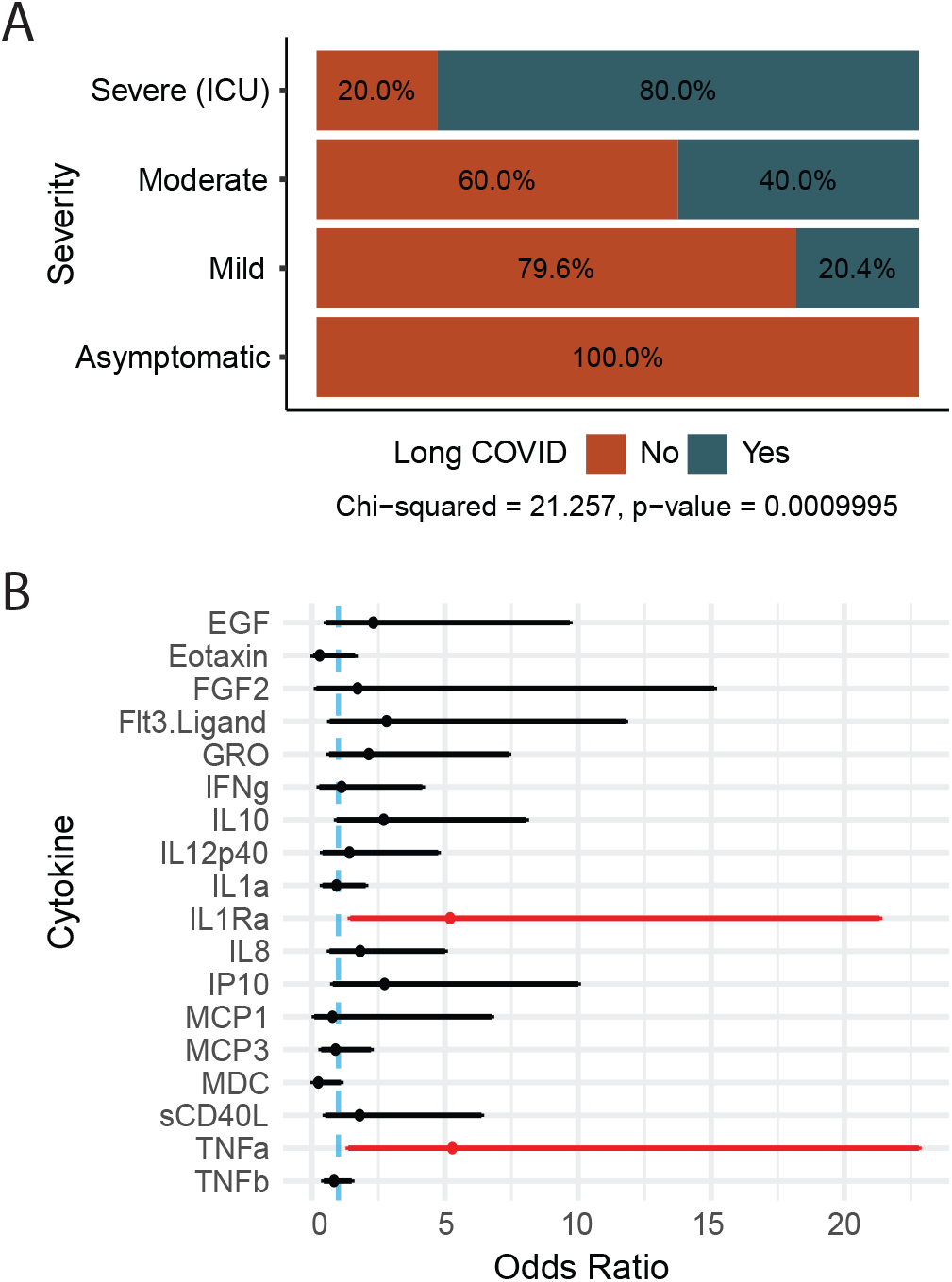
IL1Ra and TNFa are biomarkers for high risk of long COVID. (A) Prevalence of long COVID among severity groups. (B) Forest plots depicting odds ratios with 95% confidence intervals from multivariate logistic regression (log10 scale). Dashed blue line is odds ratio = 1, and red indicates a significant result, p < 0.05 and p_adj_ < 0.2. Cytokine levels were from the most acute time point available for each subject.

To determine if any cytokines were associated with development of long COVID, multivariate logistic regression was used to correct for age, sex, and days since symptom onset, and results were adjusted for multiple comparisons. We observed a ∼5-fold increase in the odds of long COVID per unit increase (on a log_10_ scale) in IL1Ra (p = 0.015, p_adj_ = 0.17) and TNFa (p = 0.019, p_adj_ = 0.17) (**Fig 3B**). Age, days since symptom onset, and sex were not significantly associated with long COVID in this analysis. A follow-up modular analysis of pairwise cytokine correlations confirmed IL1Ra and TNFa analytes did not correlate with each other, and thus represent independent signals (**Supp Fig 6**).

### Hypercytokinemia patterns are infection dependent

Another respiratory virus infection commonly associated with cytokine storm is influenza. To determine whether hypercytokinemia profiles were similar across two distinct respiratory virus infections, we utilized data from a prior study in our lab that used the same assay to identify cytokine signatures associated with influenza illness outcome (Oshansky et al. 2014) and compared the cytokine profiles across: CAH, MIS-C, Non-CAH (non-hypercytokinemia profiles in the COVID cohort), IAH (influenza associated hypercytokinemia), and Non-IAH (non-hypercytokinemia profiles in the influenza cohort). To account for the altered reference distribution, because the influenza cohort generally had higher prevalence of hypercytokinemia, we modified the classification threshold of infection associated hypercytokinemia to 20% or more of cytokines >2 standard deviations (SDs) from the acute infected mean cytokine level.

Coinciding with our prior work (Mudd et al. 2020), comparison of the average distance from the mean for a subject’s cytokine profile shows influenza induces more hypercytokinemia relative to COVID, both CAH and MIS-C (**Fig 4A**). Additionally, there are distinct patterns of elevation that can be utilized to distinguish between hypercytokinemia subtypes. IL6, GMCSF, TNFb and Age are the minimum number of factors required to classify an individual as Non-Hypercytokinemia, CAH, MIS-C, or IAH, with a misclassification rate of 2.74% (6 out of 219) (**Fig 4B**). A more comprehensive assessment of the prevalence of a cytokine as an extreme value (>2 standard deviations from the mean) for a given hypercytokinemia subtype showed IAH is the most broadly hypercytokinemic and involves a wide range of analytes (**Fig 4C**). MIS-C is the most narrow, and CAH is in between with analytes that are uniquely present as an extreme value at a higher frequency (>50%), such as MCP3 and IL9.

**Figure 4.**
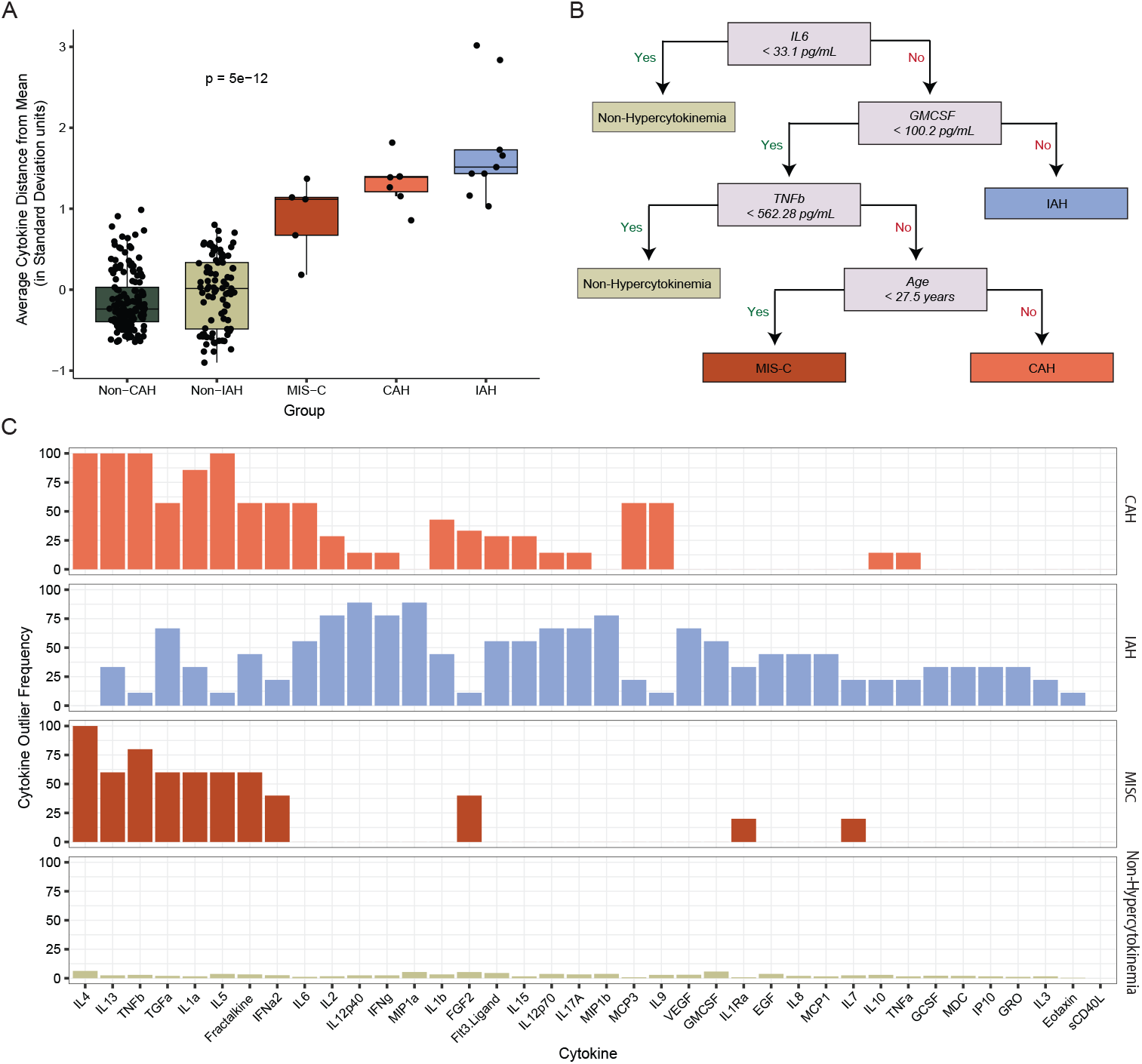
Hypercytokinemia profiles vary by respiratory virus infection. (A) Box plots of the average cytokine distance from the mean (in standard deviation units), separated and colored by hypercytokinemia group. Boxplot elements are as follows: center line, median; box limits, lower and upper quartiles; whiskers, 1.5x interquartile range; dots, individual data points.Written p-value is from a Kruskal-Wallis test across all groups. (B) Classification tree analysis to distinguish non-hypercytokinemia, CAH, IAH, and MIS-C immune profiles. (C) Bar graphs display how often a cytokine is significantly elevated (>2 standard deviations) relative to the mean, separated and colored by hypercytokinemia group.

## Discussion

This study underscores the need to reevaluate the current model of hypercytokinemia, and refine the definition of CSS as a subset within it that requires both clinical severity and broadly elevated cytokines. Amore precise, broadly applicable quantitative metric is needed, based on comprehensive analytes that can distinguish between severe hyperinflammatory disease etiologies.

These data confirmed MIS-C is a true example of CSS with nearly universally increased cytokines and high clinical severity. While CAH by definition had elevated cytokine levels, it was not associated with COVID severity. The CAH subjects in this study included 3 Mild, 3 Moderate, and 1 Severe COVID infections.Thus, although CAH can be considered an immunological feature of select COVID presentations, it is not in itself a correlate of disease severity. Including CAH as a component of their original severity group categorization did not change results, but generally increased variance, as determined by coefficient of variation and standard deviation (**Supp Fig 2-3**). This demonstrates the limitations of solely using pre-defined severity categories, as opposed to unsupervised analytical approaches, to resolve distinct immunophenotypes in response to infection, whether or not they are associated with severity. Taken together, this suggests that CAH is necessary but insufficient to classify a subject as having CSS. Including a set of minimum clinical criteria into a CSS definition would facilitate the distinction between hypercytokinemia that presents with severe clinical outcomes, such as MIS-C, and that which is not apparently clinically relevant.

We propose to modify the current paradigm of CSS, such that it is a subgroup of hypercytokinemia that is consistently associated with severe disease, and for which there are distinct types depending on disease etiology (**Fig 5**). Under this framework, CSS would appropriately include diseases such as severe sepsis and cytokine release syndrome (post CAR T cell therapy), which have high severity and clinical criteria outlined, such as that provided by the International Pediatric Sepsis Consensus Conference and the Penn Grading System, respectively (Diorio et al. 2020). Due to its lack of association with severity, CAH is a distinct form of hypercytokinemia that is associated with acute infection, but does not qualify as a subtype of CSS. Instances of hypercytokinemia that are not correlated with severe outcome, whether at a basal or acute immune state, are a distinct immunological feature that warrants further investigation and could provide valuable insight into mitigating host damage in populations with underlying inflammatory conditions, while maintaining a generally fully functional immune system.

**Figure 5.**
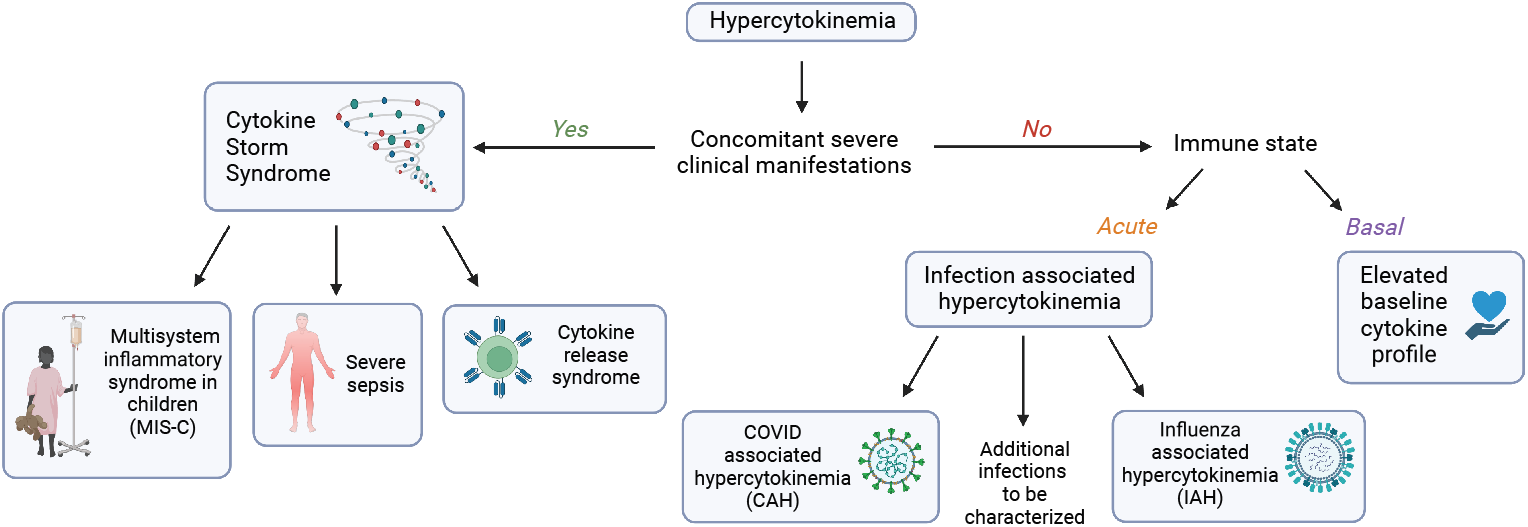
Classification of hypercytokinemia subtypes. Flow diagram of classification of hypercytokinemia based on severe clinical presentation, immune state, and disease etiology. *This image was created with BioRender.

The ideal definition of a hypercytokinemia subtype will also require quantification of the degree of elevation, and consideration of the specific analytes involved, as different hyperinflammatory diseases may have unique signatures of elevated cytokines. In comparison, the current binary classification of CSS is based solely on overcoming an undefined threshold of subjective analytes and is insufficient to characterize inflammatory profiles. It is more appropriate to assess whether a given illness outcome is associated with selective inflammation of a few pertinent analytes versus a global hyperinflammatory response. In addition, a better understanding is needed of the most appropriate and the minimum number of analytes required to define subtypes of hypercytokinemia and/or CSS (e.g., MIS-C versus severe sepsis). An optimal panel would allow for discernment between: (1) Analytes that are generally elevated during infection and, thus, would be consistently increased in all infected versus uninfected subjects. (2) Those that are uniquely associated with a given illness outcome, such as hospitalized versus non-hospitalized. (3) Those that are distinctly increased during severe hyperinflammatory states, such as MIS-C. Indeed, this study and prior work suggests that there are unique cytokine signatures within broadly hypercytokinemic conditions. A study by Diorio et al. showed pediatric cytokine release syndrome (CRS) patients could be distinguished from intensive care unit severe sepsis patients by IFNg > 83 pg/mL or IFNg < 83 pg/mL with IL1b < 8 pg/mL with 97% accuracy (Diorio et al. 2020). Based on our data, a panel that contains analytes pertinent in respiratory virus infection should include IL1b, GCSF, IFNg, IL6, GMCSF, and TNFb, as these were the minimum number of analytes required to classify “Non-Hypercytokinemia”, “MIS-C”, “CAH”, and/or “IAH” (**Fig 2C, Fig 4B**).

In summary, these results showed unique cytokine profiles and dynamics across the COVID severity spectrum, and identified IL1Ra and TNFa as putative biomarkers for patients at high risk for long COVID. Taken together, these data suggest that specific cytokine levels, obtained at key time points, may be utilized in clinical screening to identify a subject’s likely severity trajectory and/or development of long-term sequelae. Additionally, we provide a framework to improve the definition of CSS to better characterize its role in infectious diseases and can ultimately be used to improve clinical study design, reproducibility, and therapeutic strategies.

Although this study provided valuable insight into COVID cytokine profiles, there are limitations to this study. Small sample sizes for more severe groups (Moderate, Severe, and MIS-C) prevented the use of larger, multivariate models and group associated correlation analyses. Additionally, there were limited numbers of samples collected close to symptom onset, which would have allowed for more complex multivariate long COVID or severity predictive modeling with testing and training data sets.

## Supplemental Item Legends

**Supplemental Table 1.** Cohort demographics.

|  |  | n | % |
| --- | --- | --- | --- |
| Median age (years) [range] |  | 40 [7-89] |  |
| Sex |  |  |  |
|  | Female | 117 | 69.2% |
|  | Male | 52 | 30.8% |
| Race |  |  |  |
|  | Black or African American | 32 | 18.9% |
|  | White or Caucasian | 101 | 59.8% |
|  | Other/Not reported | 30 | 17.8% |
| Ethnicity |  |  |  |
|  | Hispanic | 4 | 2.4% |
| COVID Severity |  |  |  |
|  | Asymptomatic | 8 | 4.7% |
|  | Mild | 117 | 69.2% |
|  | Moderate | 22 | 13.0% |
|  | Severe | 17 | 10.1% |
| MIS-C |  | 5 | 3.0% |
| Comorbidities |  |  |  |
|  | Heart disease | 30 | 17.8% |
|  | Chronic respiratory disease | 26 | 15.4% |
|  | Current or past smoker | 25 | 14.8% |
|  | Obesity | 56 | 33.1% |
|  | Diabetes | 21 | 12.4% |
|  | Immunosuppressed | 4 | 2.4% |
|  | Long COVID (symptoms > 4 weeks) | 38 | 22.5% |
| Treatments |  |  |  |
|  | Hospitalized | 32 | 18.9% |
|  | ICU | 11 | 6.5% |
|  | Steroids | 32 | 18.9% |
|  | COVID-directed anti-viral | 13 | 7.7% |
|  | Immunomodulator | 3 | 1.8% |
| COVID vaccine breakthrough |  | 1 | 0.6% |
| Median hospital LOS, all subjects (days) |  | 1.6 |  |
| Median hospital LOS, of hospitalized subjects (days) |  | 4.6 |  |

**Supplemental Figure 1.**
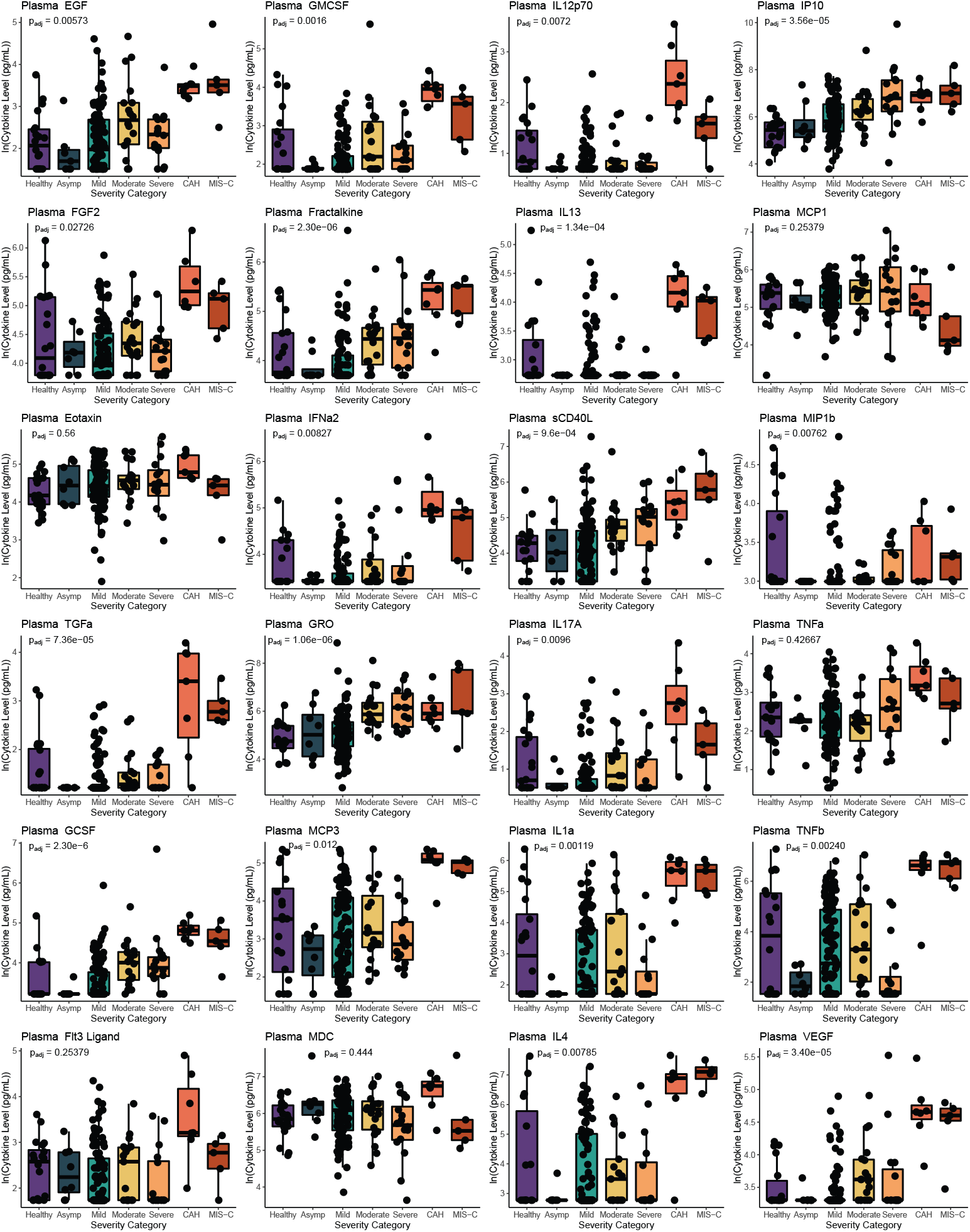
Box plots of acute cytokine levels by COVID severity group. Boxplot elements are as follows: center line, median; box limits, lower and upper quartiles; whiskers, 1.5x interquartile range; dots, individual data points. P-values are from a Kruskal-Wallis test for all groups, except CAH since its definition is based on statistical outlier calculations. Results were adjusted for multiple comparisons by controlling the FDR.

**Supplemental Figure 2.**
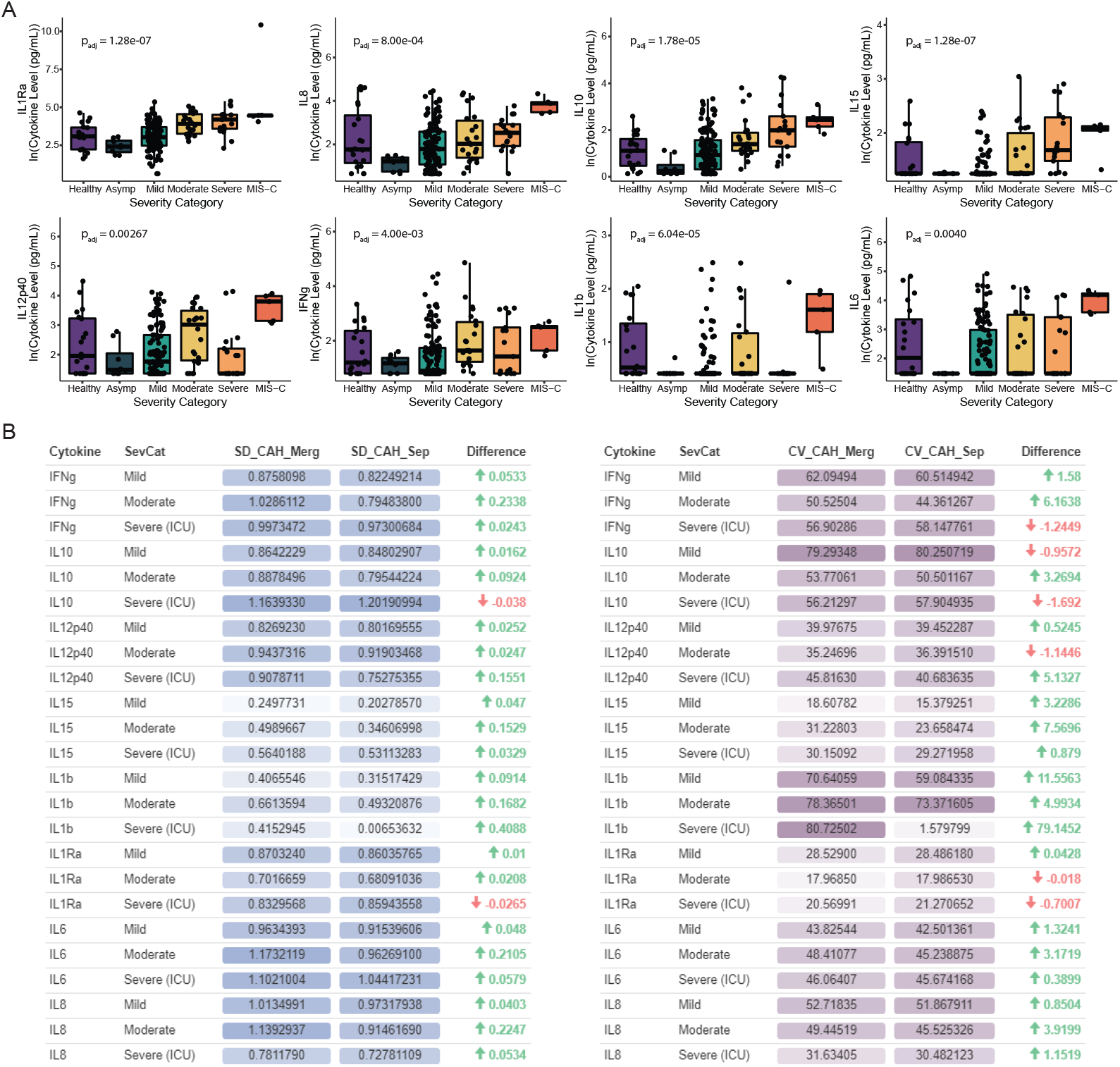
Merging CAH into severity group does not change results but increases variance. (A) Figure 2B box plots of acute cytokine levels by COVID severity group with CAH merged. P-values are from a Kruskal-Wallis test for all groups, after subjects designated as having “CAH” were merged with their original severity category. Boxplot elements are as follows: center line, median; box limits, lower and upper quartiles; whiskers, 1.5x interquartile range; dots, individual data points. Results were adjusted for multiple comparisons by controlling the FDR. (B) Left: Table of the difference in standard deviation (on a log scale) before (SD_CAH_Sep) and after (SD_CAH_Merg) merging CAH subjects with their severity group. Right: Table of the difference in coefficient of variation before (CV_CAH_Sep) and after (CV_CAH_Merg) merging CAH subjects with their severity group. Cells reporting raw values (SD_CAH_Merg, SD_CAH_Sep, CV_CAH_Merg, CV_CAH_Sep) are colored by SD or CV values to aid in visualization of differences, such as the difference in CV for IL1b in the Severe group.

**Supplemental Figure 3.**
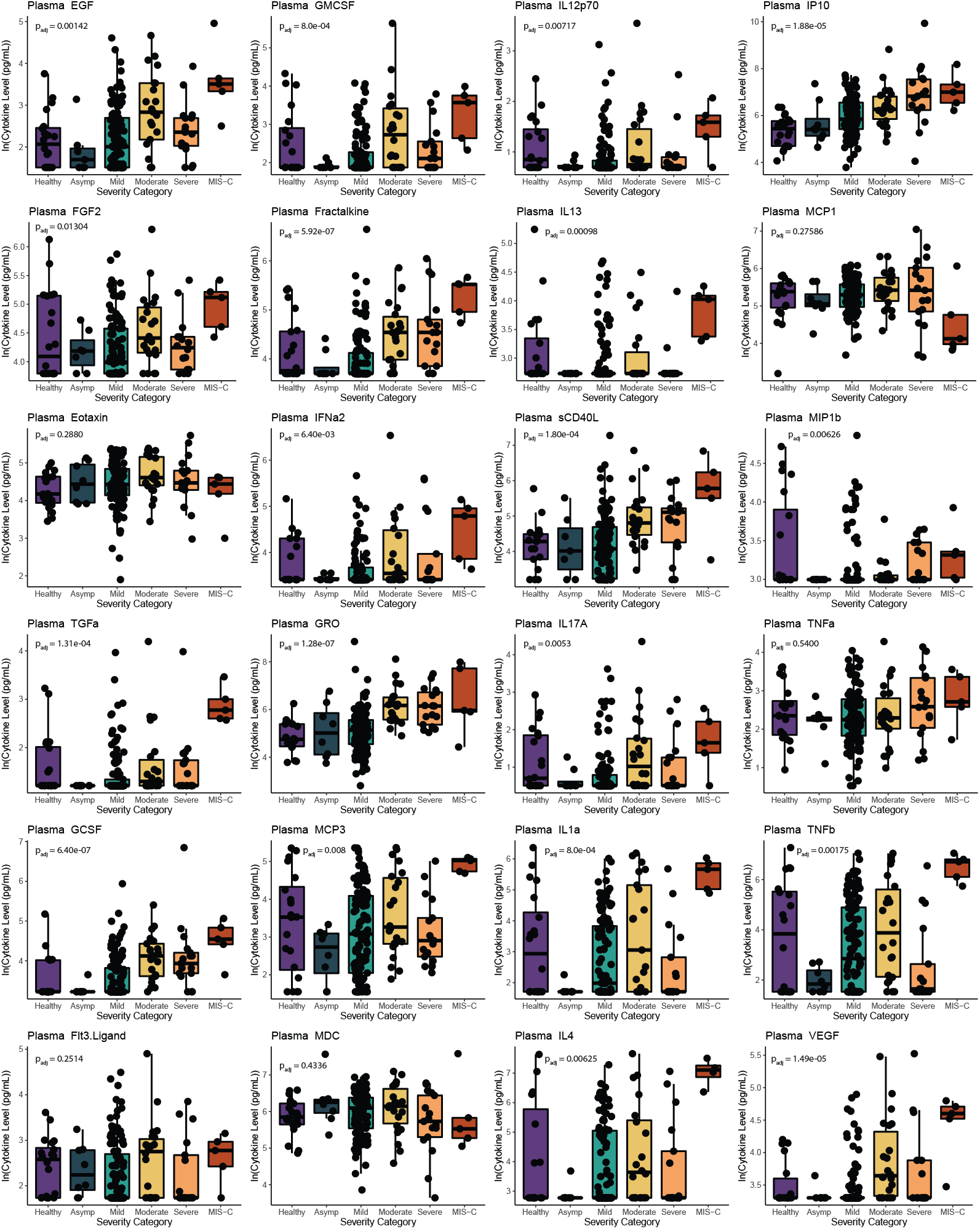
Box plots of acute cytokine levels by COVID severity group with CAH merged. P-values are from a Kruskal-Wallis test for all groups, after subjects designated as having “CAH” were merged with their original severity category. Boxplot elements are as follows: center line, median; box limits, lower and upper quartiles; whiskers, 1.5x interquartile range; dots, individual data points. Results were adjusted for multiple comparisons by controlling the FDR.

**Supplemental Figure 4.**
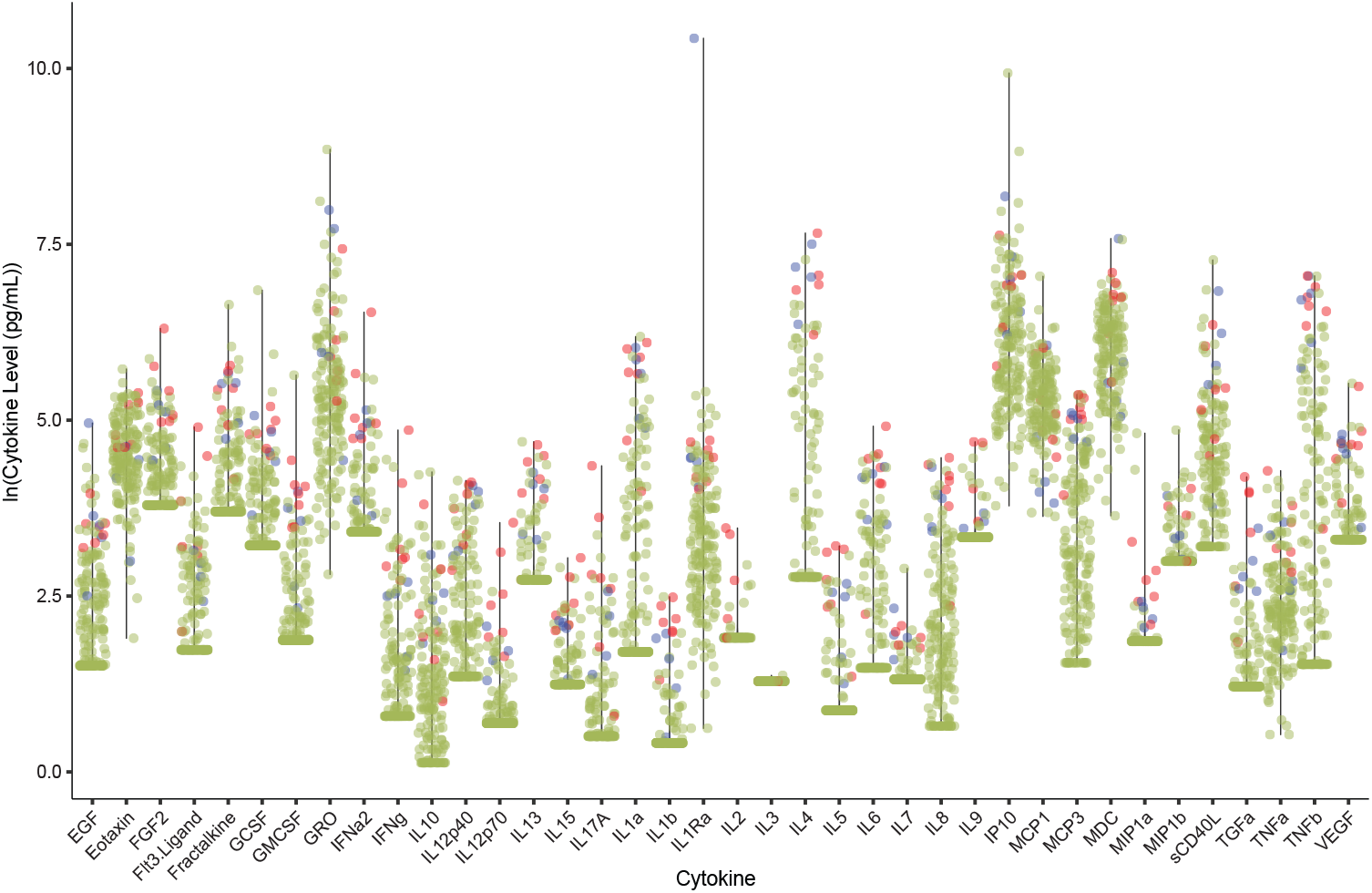
Distribution of cytokine levels in acute COVID patients. Data from the most acute time point for COVID patients (< 6 days from enrollment). CAH are shown in red, MIS-C are shown in blue, and all remaining subjects are shown in green. Lines represent the dynamic range of the data for a given analyte.

**Supplemental Figure 5.**
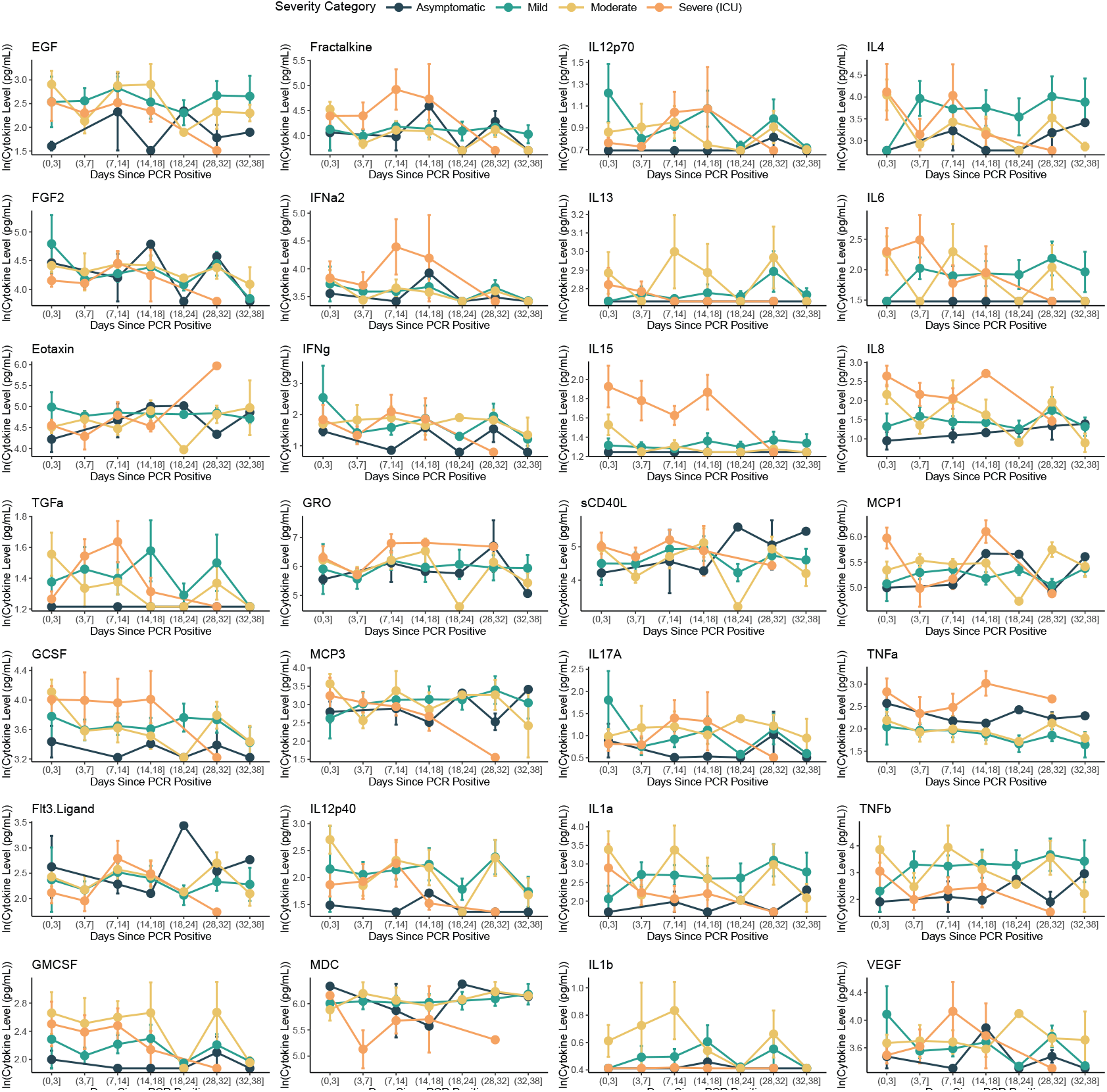
COVID cytokine dynamics. Plasma cytokine levels throughout the course of infection and colored by disease severity (n_Total_ = 151, n_Unique_ = 60). Points represent the mean and lines represent standard error.

**Supplemental Figure 6.**
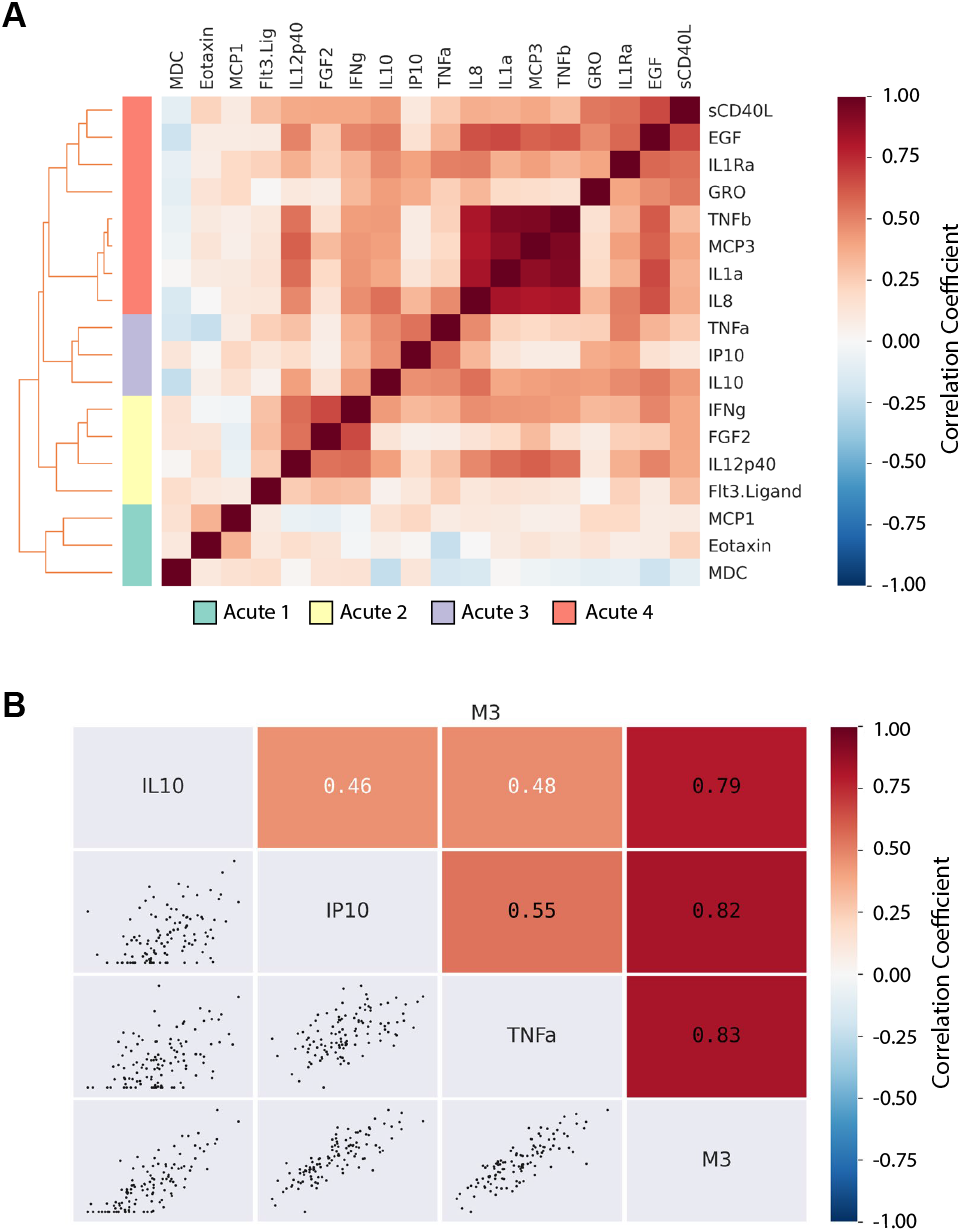
Modules of co-correlated cytokines. (A) Absolute, log-transformed cytokine concentrations were analyzed with CytoMod (Liel et al 2019). K-means clustering was utilized to identify modules of correlation, with the gap statistic identifying k=4 as optimal. (B) Correlations from cytokines in Acute Module 3 are plotted as scatterplots. In all cases, correlation coefficients represent Pearson coefficients.

## Acknowledgements

We thank the participants and clinical study support staff in each of the cohorts. This work was funded by: American Lebanese Syrian Associated Charities (ALSAC) at St. Jude Children’s Research Hospital, the Children’s Foundation Research Institute at Le Bonheur Children’s Hospital, the Collaborative Research Network at the University of Tennessee Health Science Center, the National Institute of Allergy and Infectious Diseases, US National Institutes of Health, under HHS contract HHSN272201400006C for the St. Jude Center of Excellence for Influenza Research and Surveillance (SJ CEIRS) (H.S., A.G. P.G.T.), HHS contract 75N93019C00052 for the Center for Influenza Vaccine Research for High Risk Populations (CIVR-HRP) of the Collaborative Influenza Vaccine Innovation Centers (J.W., P.G.T.), HHS contract 75N93021C00016 for the St. Jude Centers of Excellence for Influenza Research and Response (SJ CEIRR) (P.G.T.), and 5U01AI150747-03 (P.G.T.). The content is solely the responsibility of the authors and does not necessarily represent the official views of the National Institutes of Health.

## Conflicts of Interest

A.S., J.C.C., and P.G.T. have patent application(s) related to the treatment of COVID. Le Bonheur Children’s Hospital has received funds from Regeneron Pharmaceuticals and Pfizer to conduct clinical trials of COVID therapeutic agents. R.T.H. has served on advisory boards for Abbott Laboratories, Roche Diagnostics, T2 Diagnostics, and MiraVista Labs. P.G.T. has consulted or received honorarium and/or travel support from Illumina, JNJ, Pfizer, and 10X. P.G.T. serves on the Scientific Advisory Board of ImmunoScape and CytoAgents. The remaining authors declare no competing interests.

## Data Availability

Source data can be obtained from the authors upon reasonable request.

## Code Availability

Analyses were conducted using R or Python, both publicly available software, as detailed in the Methods. All code is available upon request.

## Author Contributions

Conceptualization, A.S., A.G., and P.G.T.; Methodology, A.S., A.G., and P.G.T.; Investigation, A.S. and A.G.; Formal Analysis, A.S. and J.C.C.; Visualization, A.S. and J.C.C.; Writing – Original Draft, A.S.; Writing – Review & Editing, A.S., J.C.C., A.G., and P.G.T.; Resources, J.W., A.B., M.A., L.B., K.M., N.H., C.H., A.B., S.A., S.S.T., H.S., A.G., P.G.T.; Supervision, P.G.T.; Funding Acquisition, J.W., H.S., A.G., and P.G.T.

## Methods

### Study Populations

The cohort was composed of two prospective, longitudinal, non-interventional study populations from Memphis, Tennessee: St. Jude Tracking of Viral and Host Factors Associated With COVID-19 (SJTRC; NCT04362995) and Clinical, Immunological, and Virological Characterization of COVID-19 (CIVIC-19). These studies were approved by the Institutional Review Boards of St. Jude Children’s Research Hospital and the University of Tennessee Health Science Center/Le Bonheur Children’s Hospital, respectively.

Aside from healthy controls, all participants included in these analyses tested positive for SARS-CoV-2 or were diagnosed with MIS-C. SJTRC enrolled adult St. Jude Children’s Research Hospital employees during a period of mandatory surveillance for SARS-CoV-2 via RT-PCR of mid-nasal turbinate swabs (Minervina et al. 2022). Blood samples were collected at an asymptomatic baseline and after SARS-CoV-2 infection. Healthy controls are a randomly sampled subset of SJTRC baseline samples. CIVIC-19 enrolled pediatric and adult participants who tested positive for SARS-CoV-2 by RT-PCR at one of several community hospitals or testing centers. Blood samples were collected at time of enrollment, 1-, 2-, and 4-weeks post enrollment. Samples were collected from May 2020 through March 2021. Pregnant women were excluded, but no other underlying comorbidities were excluded. None of the participants were fully vaccinated against COVID, only two Moderate severity participants had received one dose of mRNA COVID vaccine at the time of sample collection.

Written informed consent was obtained from all participants. Demographics, medical history, and any COVID symptoms, effects, treatments and vaccines were collected from electronic medical records and patient questionnaires, as available. Patients with Asymptomatic or Mild COVID often did not have electronic records available. Severity scores were assigned at a convalescent time point, 28 or more days from onset of symptoms, or SARS-CoV-2 testing if asymptomatic. The severity scoring rubric used was modified from the NIH therapeutic treatment guidelines case definition (https://www.covid19treatmentguidelines.nih.gov/overview/clinical-spectrum/) with a more stringent delineation between mild and severe cases: Asymptomatic (no symptoms reported from baseline); Mild (any cough, fever, diarrhea, vomiting, headache, loss of taste or smell, sore throat, myalgias, fatigue, lymphadenopathy or malaise *without* shortness of breath or hospitalization); Moderate (at least one of the following: oxygen saturation <92% on room air, shortness of breath, new oxygen requirement, hospitalization for COVID); Severe (at least one of the following: invasive or positive-pressure oxygen requirement, acute respiratory distress syndrome (ARDS), extra-pulmonary organ effects [acute renal injury, liver enzyme or function abnormality, coagulopathy, thrombotic event]). MIS-C was considered separate from acute COVID, and was defined as hospitalized participants <21 years of age with positive SARS-CoV-2 antibodies at time of presentation and clinical diagnosis of MIS-C given by treating physicians. Data were managed using the REDCap electronic data capture tools hosted at St. Jude and UTHSC (Harris et al. 2019).

### Specimens

Blood samples were collected by peripheral venous puncture into sodium citrate cell preparation tubes (BD Cat. #362761), centrifuged within 24 hours according to manufacturer’s recommendations to separate plasma from cellular components, and plasma was stored at - 80°C.

### Cytokine Measurement

Plasma cytokines were measured using the Luminex MAP system with the Milliplex HCYTOMAG-60K assay, and samples were processed according to the kit manual. A subject was classified as having “COVID associated hypercytokinemia” (CAH) if 30% or more of their cytokines were >2 standard deviations (SDs) from the mean cytokine level. This was calculated using all cytokine levels from the most acute time point for all SARS-CoV-2 infected patients. Thus, the “mean” in reference here represents the acute infected population mean.

### Statistical Analysis

Analyses were performed with R or Python. Prior to statistical analysis, analytes were removed from the dataset if less than 25-50% of their values were above or below the lower and upper limits of detection, respectively. This value was selected based on total sample size and the prevalence of independent and dependent variables of interest.

R analyses include comparisons across severity groups using Mann-Whitney U (2 groups), Kruskal-Wallis (3 or more groups), or chi-squared tests, depending on whether the variable of interest was numeric or categorical. Dimension reduction techniques included principal component analysis, and any missing values were replaced with the median (based on all subjects) of that cytokine level.

Python was used for correlation analyses with CytoMod (Cohen et al. 2019). For these analyses, we only analyzed samples obtained during the acute time point, and only considered cytokines for which we had sufficient observations above the minimum limit of detection (i.e., EGF, FGF2, Eotaxin, Flt3.Ligand, IFNg, GRO, IL10, MCP3, IL12p40, MDC, sCD40L, IL1Ra, IL1a, IL8, IP10, MCP1, TNFa, TNFb). Cytokines were log transformed, and co-correlating cytokines were grouped into distinct modules utilizing k-means, with the gap statistic identifying an optimal k of 4. Logistic regression for log-transformed cytokine associations with Long Covid was conducted in R with the *glm* function, with log-transformed age (in years), sex, and days since symptom onset included as covariates.

When appropriate and indicated, p-values were adjusted for multiple comparisons using the Benjamini-Hochberg method for controlling the false discovery rate (Benjamini and Hochberg 1995). For cytokine analyses, all analyte values were included to determine “CAH” designation and in the Principal Component Analysis. Prior to comparative statistical analyses, cytokine analytes were removed from the dataset, and thus subsequent statistical testing, if less than 25-50% of their values were above or below the lower and upper limits of detection, respectively. This value was selected based on total sample size and the prevalence of independent and dependent variables of interest.

## References

Benjamini, Yoav, and Yosef Hochberg. 1995. “Controlling the False Discovery Rate: A Practical and Powerful Approach to Multiple Testing.” Journal of the Royal Statistical Society. Series B (Methodological) 57 (1): 289–300.

Cohen, Liel, Andrew Fiore-Gartland, Adrienne G. Randolph, Angela Panoskaltsis-Mortari, Sook-San Wong, Jacqui Ralston, Timothy Wood, et al. 2019. “A Modular Cytokine Analysis Method Reveals Novel Associations With Clinical Phenotypes and Identifies Sets of Co-Signaling Cytokines Across Influenza Natural Infection Cohorts and Healthy Controls.” Frontiers in Immunology 10. https://www.frontiersin.org/articles/10.3389/fimmu.2019.01338.

Diorio, Caroline, Pamela A. Shaw, Edward Pequignot, Alena Orlenko, Fang Chen, Richard Aplenc, David M. Barrett, et al. 2020. “Diagnostic Biomarkers to Differentiate Sepsis from Cytokine Release Syndrome in Critically Ill Children.” Blood Advances 4 (20): 5174–83. https://doi.org/10.1182/bloodadvances.2020002592.

Fajgenbaum, David C., and Carl H. June. 2020. “Cytokine Storm.” New England Journal of Medicine 383 (23): 2255–73. https://doi.org/10.1056/NEJMra2026131.

Gallo Marin Benjamin, Ghazal Aghagoli, Katya Lavine, Lanbo Yang, Emily J. Siff, Silvia S. Chiang, Thais P. Salazar-Mather, et al. 2021. “Predictors of COVID-19 Severity: A Literature Review.” Reviews in Medical Virology 31 (1): e2146. https://doi.org/10.1002/rmv.2146.

Harris, Paul A., Robert Taylor, Brenda L. Minor, Veida Elliott, Michelle Fernandez, Lindsay O’Neal, Laura McLeod, et al. 2019. “The REDCap Consortium: Building an International Community of Software Platform Partners.” Journal of Biomedical Informatics 95 (July): 103208. https://doi.org/10.1016/j.jbi.2019.103208.

Leisman, Daniel E., Lukas Ronner, Rachel Pinotti, Matthew D. Taylor, Pratik Sinha, Carolyn S. Calfee, Alexandre V. Hirayama, et al. 2020. “Cytokine Elevation in Severe and Critical COVID-19: A Rapid Systematic Review, Meta-Analysis, and Comparison with Other Inflammatory Syndromes.” The Lancet Respiratory Medicine 8 (12): 1233–44. https://doi.org/10.1016/S2213-2600(20)30404-5.

Michelen, Melina, Lakshmi Manoharan, Natalie Elkheir, Vincent Cheng, Andrew Dagens, Claire Hastie, Margaret O’Hara, et al. 2021. “Characterising Long COVID: A Living Systematic Review.” BMJ Global Health 6 (9): e005427. https://doi.org/10.1136/bmjgh-2021-005427.

Minervina, Anastasia A., Mikhail V. Pogorelyy, Allison M. Kirk, Jeremy Chase Crawford, E. Kaitlynn Allen, Ching-Heng Chou, Robert C. Mettelman, et al. 2022. “SARS-CoV-2 Antigen Exposure History Shapes Phenotypes and Specificity of Memory CD8+ T Cells.” Nature Immunology 23 (5): 781–90. https://doi.org/10.1038/s41590-022-01184-4.

Mudd, Philip A., Jeremy Chase Crawford, Jackson S. Turner, Aisha Souquette, Daniel Reynolds, Diane Bender, James P. Bosanquet, et al. 2020. “Distinct Inflammatory Profiles Distinguish COVID-19 from Influenza with Limited Contributions from Cytokine Storm.” Science Advances, November, eabe3024. https://doi.org/10.1126/sciadv.abe3024.

Oshansky, Christine M., Andrew J. Gartland, Sook-San Wong, Trushar Jeevan, David Wang, Philippa L. Roddam, Miguela A. Caniza, et al. 2014. “Mucosal Immune Responses Predict Clinical Outcomes during Influenza Infection Independently of Age and Viral Load.” American Journal of Respiratory and Critical Care Medicine 189 (4): 449–62. https://doi.org/10.1164/rccm.201309-1616OC.

